# Structural Insights into Competitive Binding Dynamics between RALF23/33 and PCP-B in Brassicaceae Pollination

**DOI:** 10.1101/2024.12.03.626569

**Authors:** Hemal Bhalla, Karthik Sudarsanam, Ashutosh Srivastava, Subramanian Sankaranarayanan

**Affiliations:** Department of Biological Sciences and Engineering, Indian Institute of Technology Gandhinagar, Palaj, Gujarat - 382355, India

**Keywords:** Brassicaceae, FERONIA, LLG, PCP-B, RALF23, Structural modeling

## Abstract

Ensuring successful fertilization, viable offspring production, genetic isolation, and maintaining species integrity is pivotal for the survival of flowering plants. Members of Brassicaceae employ a “gatekeeping mechanism” involving interaction between stigmatic membrane-bound *Catharanthus roseus* receptor-like kinase 1-like (CrRLK1L) receptor, FERONIA, GPI anchored protein LLG2 (LORELEI-LIKE GLYCOPHOSPHATIDYLINOSITOL-ANCHORED PROTEIN 2) and autocrine secreted RALF23/33 (Rapid alkalinization factor) peptide. This binding establishes a barrier for pollen hydration by inducing ROS (Reactive Oxygen Species). Conversely, in the presence of compatible pollen, paracrine-secreted cysteine-rich peptides such as PCP-Bγ compete with RALF23/33 for binding to the FERONIA-LLG2 complex, thus reducing ROS levels, ensuring successful pollen hydration and germination. Despite its crucial role, the structural basis of this competitive binding dynamics remains elusive owing to the lack of structural data and the inherent flexibility of these peptides. Using structural modeling, molecular docking, and simulations, this study reveals that PCP-Bγ binds to the same negatively charged pocket in the FERONIA-LLG2 complex as RALF23, displacing and interrupting the heterodimerized structure, thus reducing ROS levels to promote pollination. Our study unveils the experimental data-based predicted models, competitive binding dynamics, and mechanism behind this “gatekeeping mechanism,” shedding light on the molecular mechanism underlying this pollen hydration barrier in Brassicaceae.

## Introduction

Flowering plants rely on precise communication between pollen and pistil to distinguish between compatible/desired and incompatible/unwanted pollen, ensuring successful fertilization, species integrity, and genetic isolation. In Brassicaceae, this “gatekeeping mechanism” involves the CrRLK1L (*Catharanthus roseus* receptor-like kinase 1-like) receptor, FERONIA, and GPI-anchored LLG 2 (LORELEI-LIKE GLYCOSYLPHOSPHATIDYLINOSITOL-ANCHORED PROTEIN 2) which acts as “gatekeeper” by perceiving autocrine secreted stigmatic peptides RALF23/33 (Rapid Alkalinization Factor). This interaction induces ROS (Reactive Oxygen Species) production in stigmatic papilla cells through the RAC/ROP-RBOHD (Rho-like GTPases-Respiratory burst oxidase homolog D) mediated pathway, inhibiting hydration of incompatible/unwanted pollen. In contrast, compatible pollen releases PCP-B peptides, which compete with RALF23/33 for binding to the FERONIA-LLG2 complex, reducing ROS levels and allowing successful pollen hydration (Fig. 1A) (Huang et al., 2023; Liu et al., 2021). Despite the dynamic interaction between RALF23/33 and PCP-Bγ, RALF23 generally exhibits a higher binding affinity for the FERONIA-LLG2 complex, suggesting that other regulatory mechanisms also influence the process (Liu et al., 2021; Zhou et al., 2021).

**Fig. 1.**
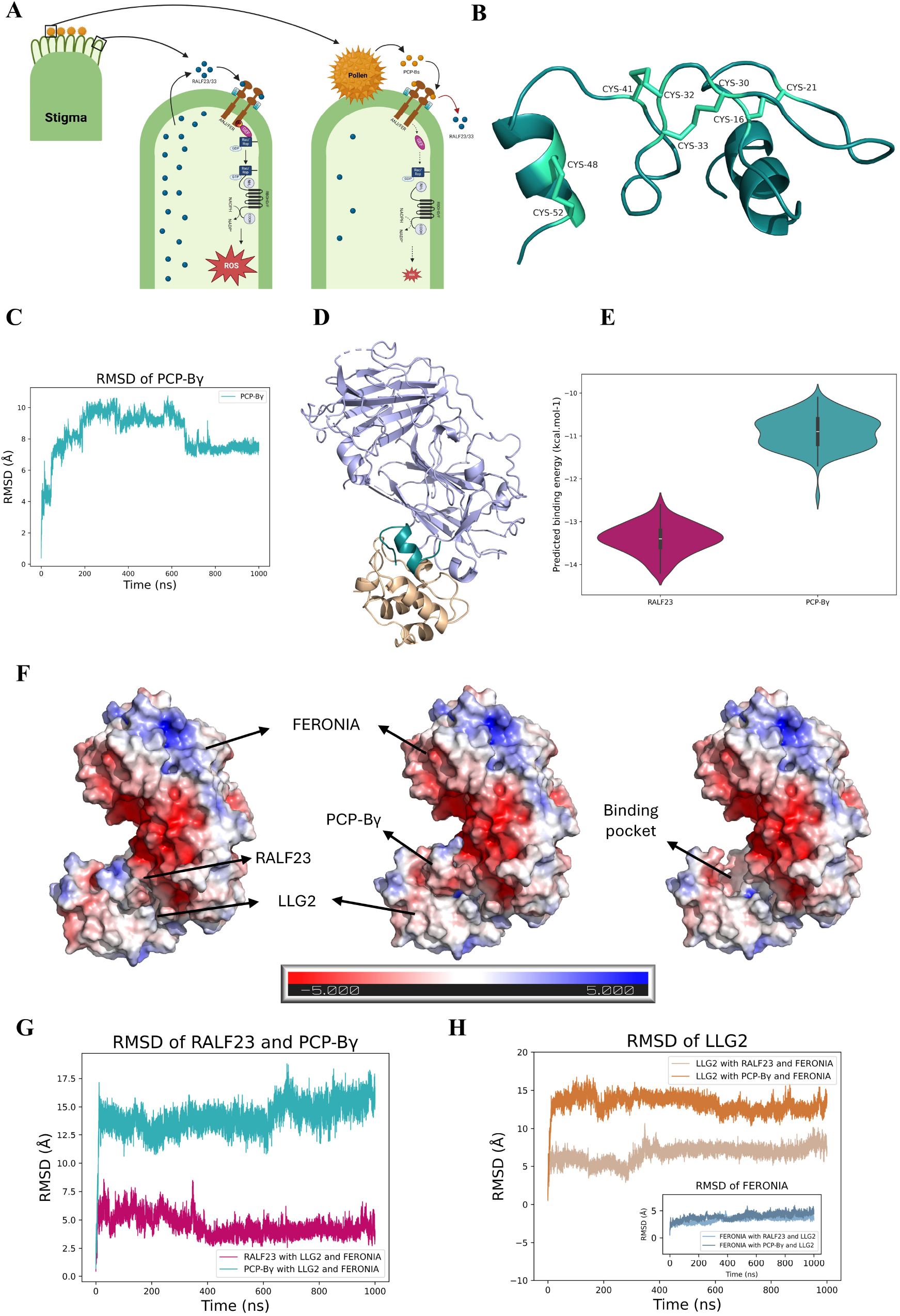
Structural insights into competitive binding dynamics between PCP-Bγ and RALF23 to FERONIA-LLG2 complex. (A) Schematic diagram showing competitive binding dynamics between PCP-Bγ (Pollen Coat Protein -Bγ) and RALF23 (Rapid Alkalisation Factor 23) to stigmatic membrane-associated FERONIA-LLG2 (LORELEI-LIKE GLYCOSYLPHOSPHATIDYLINOSITOL-ANCHORED PROTEIN 2) co-receptors. (B) Representative model of PCP-Bγ predicted using Rosetta abinitio protocol highlighting disulfide linkages. (C) Root Mean Square Deviation (RMSD) of C-α atoms of PCP-Bγ across 1000 ns MD simulation trajectory. (D) Representative FERONIA-LLG2-PCP-Bγ model predicted by HADDOCK 2.4. (Light blue: FERONIA, Cyan: PCP-Bγ, and Brown: LLG2) (E) Predicted binding affinities of RALF23 and PCP-Bγ in complex with FERONIA-LLG2 by utilizing the last 100 frames from 1000 ns simulation trajectory. (F) Representative structure of FERONIA-LLG2-RALF23 (left) and FERONIA-LLG2-PCP-Bγ (middle) and FERONIA-LLG2 (showing the binding pocket) (right), colored based on electrostatic potential as calculated by APBS. Red color indicates negative potential, and blue indicates positive potential. The darker the color, the more electrostatic potential. (G) Root mean square deviation (RMSD) of C-α atoms of RALF23 and PCP-Bγ in FERONIA-LLG2-RALF23/PCP-Bγ complex across 1000 ns MD simulation trajectory. (H) Root mean square deviation (RMSD) of C-α atoms of LLG2 in FERONIA-LLG2-RALF23/PCP-Bγ complex across 1000 ns MD simulation trajectory. The inset figure shows the RMSD of C-α atoms of FERONIA along the trajectory.

RALF23’s role is not limited to pollen hydration. This peptide is also known to induce heterodimerization of FERONIA and LLG2, contributing to a variety of plant physiological processes. Previous studies have shown that RALF23-mediated FERONIA-LLG2 heterodimerization regulates root growth (Haruta et al., 2014), enhances immunity (Stegmann et al., 2017), and mediates pollen acceptance (Liu et al., 2021). Notably, the structure of RALF23 has been partially resolved, and crystallographic studies have successfully captured the N-terminal region of RALF23 in complex with the ectodomains of FERONIA and LLG2 (Supplementary Fig. S2A) (Xiao et al., 2019). This N-terminal region is highly conserved and plays a crucial role in the biochemical recognition of RALF23 by its cognate receptors (Pearce et al., 2010).

Cysteine-rich proteins (CRPs) like PCP-Bγ, which are key regulators of pollen hydration, have been identified in various species, including Arabidopsis (Liu et al., 2021; Wang et al., 2017). These peptides are characterized by functionally important cysteine residues that form disulfide bonds, which are essential for maintaining the structural integrity of these molecules. The functional importance of these cysteine residues is highlighted by experimental evidence showing a loss of activity in modified PCP-Bγ peptides where cysteines were altered (Liu et al., 2021). These peptides contain eight conserved cysteines, which form four disulfide bonds, maintaining their functional conformation during interactions with the FERONIA-LLG2 receptor complex (Liu et al., 2021; Wang et al., 2017). Although the crystal structure of these peptides has not yet been determined, structural models have been predicted (Supplementary Fig. S1A) (Jumper et al., 2021; Wang et al., 2017).

Despite these insights, the structural basis for the competitive binding dynamics between RALF23/33 and PCP-Bγ remains elusive. This is primarily due to the inherent flexibility of these peptides and the lack of high-resolution experimental data on their interaction with the FERONIA-LLG2 complex (Xiao et al., 2019). Understanding the structural details of this competitive binding is critical for unraveling the molecular mechanisms underlying pollen recognition and hydration.

In this study, we employ a combination of structural modeling, molecular docking, and molecular dynamics simulations to explore the competitive binding dynamics of RALF23/33 and PCP-Bγ with the FERONIA-LLG2 complex. By leveraging these advanced computational techniques, we aim to provide new insights into the molecular basis of this “checkpoint” mechanism, offering a deeper understanding of how pollen recognition and hydration are regulated in flowering plants.

## Results and Discussion

To investigate the binding dynamics between RALF23/PCP-Bγ with FERONIA-LLG2, the structure of PCP-Bγ was modeled *ab initio* and docked with FERONIA-LLG2 crystal structure using Haddock 2.4 by providing experimental restraints (Dominguez et al., 2003; van Zundert et al., 2016). Structure and sequence information of RALF23/PCP-Bγ and FERONIA-LLG2 were retrieved from UniProt and Protein Data Bank (Berman et al., 2000; The UniProt Consortium et al., 2024). The crystal structure of FERONIA-LLG2-RALF23 (PDB ID: 6A5E) had only the N-terminal of RALF23 resolved, possibly due to the high flexibility of the C-terminal region, while certain regions of FERONIA were also missing (Supplementary Fig. S2A) (Berman et al., 2000; Xiao et al., 2019). The predicted model of PCP-Bγ derived from the Alpha Fold database (AF-A8MR88) indicated the presence of three disulfide bonds between C16-33, C21-52, and C30-48 in contradiction to four bonds between C16-21, C30-33, C32-41 and C48-52 reported earlier using mass spectrometry (Supplementary Fig. S1A, Supplementary Table S1) (Liu et al., 2021). The predicted model showed high predicted aligned error (pAE) and low predicted local distance difference test scores (pLDDT), indicating low confidence in prediction (Supplementary Fig. S1A and S1B). Secondary structure prediction of PCP-Bγ indicated the presence of primarily coils with a small helix at the C-terminal region (Supplementary Fig. S1C). AlphaFold prediction was also not consistent with this. Consequently, we used ROSETTA *ab initio* structure prediction protocol utilizing disulfide bond restraints derived from mass spectrometry-based experimental data and predicted the tertiary structure of PCP-Bγ (Supplementary Table S1) (Liu et al., 2021). The predicted structure revealed four disulfide bonds and the presence of a C-terminal helix, consistent with secondary structure predictions (Fig. 1B, Supplementary Fig. S1D, S1E). The predicted models had ROSETTA scores ranging from 1288.93 to - 61.953. Models (n = 5702) with negative ROSETTA scores were used for further analysis with RMSD (Root mean square deviation) from the highest scoring structure ranging from 0 to 17.5 Å. Most models analyzed showed ROSETTA scores in the range of 0 to -50 and RMSDs between 5 to 15 Å (Supplementary Fig. S1F). The predicted models were clustered using RMSD. The largest cluster contained 1802 models and was chosen for further analysis (Supplementary Table S2). The representative model from this cluster was chosen based on the score and relative RMSD from other models (Supplementary Table S3, Supplementary Fig. S1D). Molecular dynamics simulations of the representative PCP-Bγ model (Model 2) indicated varying RMSD ranging from 0 to 10 Å for the initial 600ns and then showed a stable pattern over the next 400ns ranging from 7 to 8 Å (Fig. 1C). The Root mean square fluctuation (RMSF) revealed relatively lower flexibility in the residues from 0 to 40 whereas higher flexibility was observed in the residues 41 to 54 (Supplementary Fig. S1G). The low flexibility regions involved the residues around disulfide bonds. This could be due to the conformational constraints caused by disulfide bonds. The radius of gyration ranged between 10 to 13 Å, indicating a stable and compact model (Supplementary Fig. S1H).

Structural homology analysis using matchmaker-Chimera 1.16 (Meng et al., 2023) revealed a similarity between the C-terminal helical region of PCP-Bγ (R40 to D54) and the N-terminal RALF23 structure (R4 to C18) crystal structure (Supplementary Fig. S2B). Structurally homologous C-terminal helical region of PCP-Bγ was docked with FERONIA-LLG2 crystal structure using ambiguous restraints (Supplementary Table S4) based on RALF23 interaction with the complex (Xiao et al., 2019). The top 6 clusters based on haddock score ranged from -82.303 to -58.282 (Supplementary Table S5, Supplementary Fig. S2C). Cluster 2 was used for further analysis based on a relatively high haddock score and lower RMSD (Fig. 1D). Structural comparison between FERONIA-LLG2-RALF23 crystal structure (PDB ID 6A5E) and FERONIA-LLG2-PCP-Bγ predicted model showed no deviation in FERONIA and LLG2 during the docking protocol while RALF23 and PCP-Bγ showed binding in similar regions (Supplementary Fig. S3A). Binding energy analysis of the last 100 frames of FERONIA-LLG2-RALF23 and FERONIA-LLG2-PCP-Bγ simulation trajectories predicted slightly lower binding energy for FERONIA-LLG2-RALF23 complex in comparison to FERONIA-LLG2-PCP-Bγ, consistent with the microscale thermophoresis analysis data (Fig. 1E) (Liu et al., 2021). Similarly, binding energy predicted using PRODIGY (Vangone and Bonvin, 2015) indicated slightly lower binding energy for the FERONIA-LLG2-RALF23 complex in comparison to the FERONIA-LLG2-PCP-Bγ complex (Supplementary Fig. S3B, Supplementary Table S6). This suggests that, *in vivo*, higher concentrations of PCP-Bγ might balance its lower binding affinity, or it may interfere with the FERONIA-LLG2 interaction, which is known to be mediated by RALF23 and is integral for ROS production (Xiao et al., 2019). Molecular surface electrostatics showed that the binding pocket where RALF23 and PCP-Bγ are known to bind is negatively charged, emphasizing the importance of positively charged residues in the helices for stable and more robust binding in the pocket (Fig. 1F). While the number of positively charged residues remains same in both the interacting helices, PCP-Bγ contains some negatively charged residues, which can be implicated in slightly higher binding energy in the negatively charged pocket due to electrostatic repulsion (Supplementary Fig. S3C). Residue-level interaction analysis revealed various polar contacts between the C terminal helix of PCP-Bγ and the FERONIA-LLG2 complex (Supplementary Fig. S3D, Supplementary Table S7). However, the number of polar contacts was higher between N-terminal RALF23 and LLG2, possibly contributing to the observed higher binding affinity of RALF23 in comparison to PCP-Bγ with FERONIA-LLG2 (Supplementary Fig. S3E, Supplementary Table S8). Interestingly, no direct contact between RALF23 and FERONIA was observed, indicating that the C terminal of RALF23 is likely responsible for this interaction (Supplementary Fig. S3E, Supplementary Table S8) (Xiao et al., 2019). Molecular dynamic simulation for FERONIA-LLG2-RALF23/PCP-Bγ complex indicated RMSD for RALF23 ranging from 2.5 to 7.5 Å, while for PCP-Bγ it ranged between 12.5 to 17.5 Å (Fig. 1G). Although both peptides remained stable within the negatively charged binding pocket, the lower RMSD of RALF23 supported its predicted higher binding affinity with the FERONIA-LLG2 complex. FERONIA remained stable throughout the 1000 ns simulation, while LLG2 destabilized and moved away from the complex in the presence of PCP-Bγ, suggesting that PCP-Bγ may disrupt the FERONIA-LLG2 interaction (Fig. 1H). Root mean square fluctuation of C-α atoms indicated higher fluctuation in LLG2 atoms while in complex with PCP-Bγcompared to RALF23 (Supplementary Fig. S4A, S4B). These observations support the notion that PCP-Bγ competes with RALF23 for the negatively charged binding pocket and leads to interruption of the FERONIA-LLG2 complex, which is nucleated by RALF23, thus reducing ROS levels.

In summary, we have successfully predicted a previously unknown model for PCP-Bγ based on available experimental data and provided a detailed analysis of its competitive interaction with RALF23 and FERONIA-LLG2 through molecular docking and simulation. These insights into the molecular mechanism acting as a “checkpoint” in controlling pollen hydration and germination were achieved by integrating experimental restraints into structural predictions and docking studies (Supplementary Fig. S5). As indicated in previous studies, our model shows displacement through competitive binding and higher binding energy of PCP-Bγ compared to RALF23 with FERONIA-LLG2. Our models complement the previous experimental findings (Liu et al., 2021) and highlight the potential role of PCP-Bγ in interrupting the FERONIA-LLG2 complex. Future experiments, including mutation studies, can further elucidate key residues in these interactions. Advanced protein structure determination methods will provide greater clarity on these binding dynamics in the future. Nevertheless, our study offers valuable insights into the mechanism by which PCP-Bγ interacts with the FERONIA-LLG2 complex, displacing RALF23 and disrupting the FERONIA-LLG2 interaction, ultimately leading to a reduction in ROS levels. Furthermore, this hybrid pipeline, which integrates experimental restraints, can be effectively applied to model various peptide and protein structures and analyze their binding dynamics.

## Acknowledgments

We acknowledge the Ministry of Education for the Prime Minister Research Fellowship to HB and IITGN for fellowship to KS. We acknowledge support from DBT for the Ramalingaswami Re-entry fellowship and IITGN start-up grants to AS and SS.

## Author contributions

HB: conceptualization, formal analysis, investigation, data curation, writing-original draft. KS: formal analysis, investigation, data curation. AS: formal analysis, investigation, writing-review and editing, supervision, funding acquisition. SS: conceptualization, investigation writing-review and editing, supervision, funding acquisition.

