## Supplemental information for "Structural Insights into Competitive Binding Dynamics between RALF23/33 and PCP-B in Brassicaceae Pollination"

**Affiliations:**

**Methods**

**Data retrieval from online repositories**

The crystal structure of FERONIA-LLG2-RALF23 was derived from the PDB repository (PDB ID: 6A5E) (Berman et al., 2000; Xiao et al., 2019). The predicted structure of PCP-Bγ was derived from the AlphaFold database (AF-A8MR88-F1-v4) (Jumper et al., 2021). Sequence information for PCP-Bγ (AT2G16535) and RALF23 (AT3G16570) was derived from NCBI (Sayers et al., 2021).

**Structure prediction and analysis of RALF23 and PCP-Bγ**

The secondary structure of PCP-Bγ was predicted using PSIPRED 4.0, available at the PSIPRED workbench (Buchan and Jones, 2019; Jones, 1999). The tertiary structure of PCP-Bγ was predicted using ROSETTA abinitio protocol by providing the disulfide bond restraints (fix_disulf flag) (Supplementary Table S1) (Liu et al., 2021). The number of models predicted was 10000 (nstruct flag). All other parameters were used as default for tertiary structure prediction (Rohl et al., 2004). Clustering was performed using ROSETTA Calibur based on the root mean square deviation of PCP-Bγ in the models predicted by ROSETTA ab initio protocol (Li and Ng, 2010). The representative predicted model of PCP-Bγ and the crystal structure of RALF23 were aligned using Chimera 1.16- matchmaker (Meng et al., 2023).

**Molecular docking of PCP-Bγ-FERONIA-LLG complex**

The experimentally derived structure of FERONIA-LLG2 was derived from PDB (PDB ID: 6A5E) (Xiao et al., 2019). The structure was loop-modelled and optimized for missing loops using the Rosetta remodel algorithm using 1000 trajectories and “quick and dirty” flag; all other parameters were set to default (Huang et al., 2011). The predicted model of PCP-Bγ was docked with the crystal structure using ambiguous restraints (Supplementary Table S4) in the Haddock 2.4 online server. All other parameters were used as default for docking (Dominguez et al., 2003; van Zundert et al., 2016).

**Binding and interaction analysis**

The representative docked model of FERONIA-LLG2-PCP-Bγ and the crystal structure of FERONIA-LLG2-RALF23 were used for binding energy analysis using the Prodigy online server (Vangone and Bonvin, 2015; Xue et al., 2016). The last 100 frames from each 1000 ns simulation trajectory were used for binding energy analysis using PRODIGY (Vangone and Bonvin, 2015; Xue et al., 2016). Molecular surface electrostatics was calculated for representative model and crystal structure using the Adaptive Poisson Boltzmann Solver (APBS) in PyMOL. The interactions between FERONIA-LLG2 and PCP-Bγ/RALF23 were analyzed using the ePISA online server (Krissinel and Henrick, 2005).

**Molecular Dynamics Simulation**

All-atom molecular dynamics (MD) simulations were performed for PCP-Bγ, FERONIA-RALF23-LLG2, and FERONIA-PCP-Bγ -LLG2 systems using GROMACS version 2021.3. The Amber99sb-ildn force field was applied for all simulations, and the TIP3P water model was utilized to solvate the protein complexes within a dodecahedral simulation box. To maintain physiological ionic strength, Na+, and Cl- ions were added to maintain the concentration of 0.15 M.

Energy minimization was executed via the steepest descent algorithm, with 50,000 steps or a maximum force of 500 kJ/mol/nm on any atom. Post-minimization, equilibration was carried out in two stages: first, 1 ns equilibration in the NVT ensemble, where the system temperature was regulated at 300 K using a modified Berendsen thermostat (Berendsen et al., 1984). This was followed by NPT equilibration in two sequential 1 ns steps, initially using the Berendsen barostat (Berendsen et al., 1984) and subsequently with the Parrinello-Rahman barostat (Parrinello and Rahman, 1981), both at 1 bar pressure. Throughout, the system temperature was maintained at 300 K using the modified Berendsen thermostat (V-rescale).

Finally, a production MD run of 1 microsecond was performed for each system under the NPT ensemble using the Parrinello-Rahman barostat with pressure at 1 bar and temperature at 300 K.

The structural properties of the simulated systems, including Root Mean Square Deviation (RMSD), Root Mean Square Fluctuation (RMSF), and Radius of Gyration (Rg), were analyzed using GROMACS tools gmx rms, gmx rmsf, and gmx gyrate, respectively.

**Supplementary Figure Legends:**

**Supplementary Figure S1. Structure prediction and analysis of PCP-Bγ.**

(A) Structure of PCP-Bγ predicted by Alpha Fold (Alpha Fold DB: A8MR88) highlighting disulfide linkages. The coloring indicates predicted local distance difference test scores (pLDDT).
(B) Graph showing predicted aligned error (PAE) plot of PCP-Bγ predicted by Alpha Fold (Alpha Fold DB: A8MR88).
(C) PSIPRED prediction of secondary structure in PCP-Bγ. 3-State prediction is denoted as follows: C (coils), E (beta strands), and H (Alpha helix).
(D) Top 5 scoring models of PCP-Bγ predicted using ROSETTA abinitio protocol using disulfide bond restraints.
(E) Representative model of PCP-Bγ predicted using Rosetta abinitio protocol highlighting disulfide linkages.
(F) Score versus root mean square deviation (RMSD) plot of PCP-Bγ models predicted using Rosetta abinitio protocol.
(G) Root mean square fluctuation (RMSF) of C-α atoms of PCP-Bγ for 1000 ns MD simulation.
(H) Radius of gyration of C-α atoms of PCP-Bγ for 1000 ns MD simulation.

**Supplementary Figure S2. Structural alignment and molecular docking of FERONIA-LLG2-PCP-Bγ complex.**

(A) Crystal structure of FERONIA-LLG2-RALF23 complex. (PDB ID: 6A5E) (Light blue: FERONIA, Pink: RALF23, and Brown: LLG2)
(B) Structural comparison of RALF23 (pink) crystal structure and predicted PCP-Bγ (cyan) model using matchmaker- Chimera1.16
(C) Superimposed FERONIA-LLG2-PCP-Bγ models of top 6 clusters predicted by HADDOCK 2.4 (Light blue: FERONIA, Cyan: PCP-Bγ and Brown: LLG2)

**Supplementary Figure S3. Residue level interaction analysis between FERONIA-LLG2-PCP-Bγ and FERONIA-LLG2-RALF23 complex.**

(A) Structural comparison between FERONIA-LLG2-RALF23 crystal structure and FERONIA-LLG2-PCP-Bγ predicted model. (Light blue: FERONIA, Cyan: PCP-Bγ, Pink: RALF23, and Brown: LLG2)
(B) Graph representing the binding energies of FERONIA-LLG2-RALF23 and FERONIA-LLG2-PCP-Bγ complex predicted using Prodigy server.
(C) Graph representing the distribution of charged residues in RALF23 (residue:4-18) and PCP-Bγ (residue:40-54).
(D) Structure of FERONIA-LLG2-PCP-Bγ showing polar contacts (left, shown as yellow dotted lines) and its magnified view (right). (Light blue: FERONIA, Cyan: PCP-Bγ, and Brown: LLG2)
(E) Structure of FERONIA-LLG2-RALF23 showing polar contacts (left, shown as yellow dotted lines) and its magnified view (right). (Light blue: FERONIA, Pink: RALF23, and Brown: LLG2)

**Supplementary Figure S4. Interaction analysis of FERONIA-LLG2-PCP-Bγ and FERONIA-LLG2-RALF23 complex by MD simulations.**

(A) Root mean square fluctuation of C-α atoms in FERONIA-LLG2-PCP-Bγ complex for 1000 ns MD simulation. The residue on the left belongs to FERONIA, the middle belongs to LLG2, and the right belongs to PCP-Bγ.
(B) Root mean square fluctuation of C-α atoms in FERONIA-LLG2-RALF23 complex for 1000 ns MD simulation. The residue on the left belongs to FERONIA, the middle belongs to LLG2, and on the right belongs to RALF23.

**Supplementary Figure S5. Schematic workflow followed in this study for structure comparison, integrative modeling, and molecular docking of the FERONIA-LLG2-PCP-Bγ complex.**

**Supplementary Figure S1.**

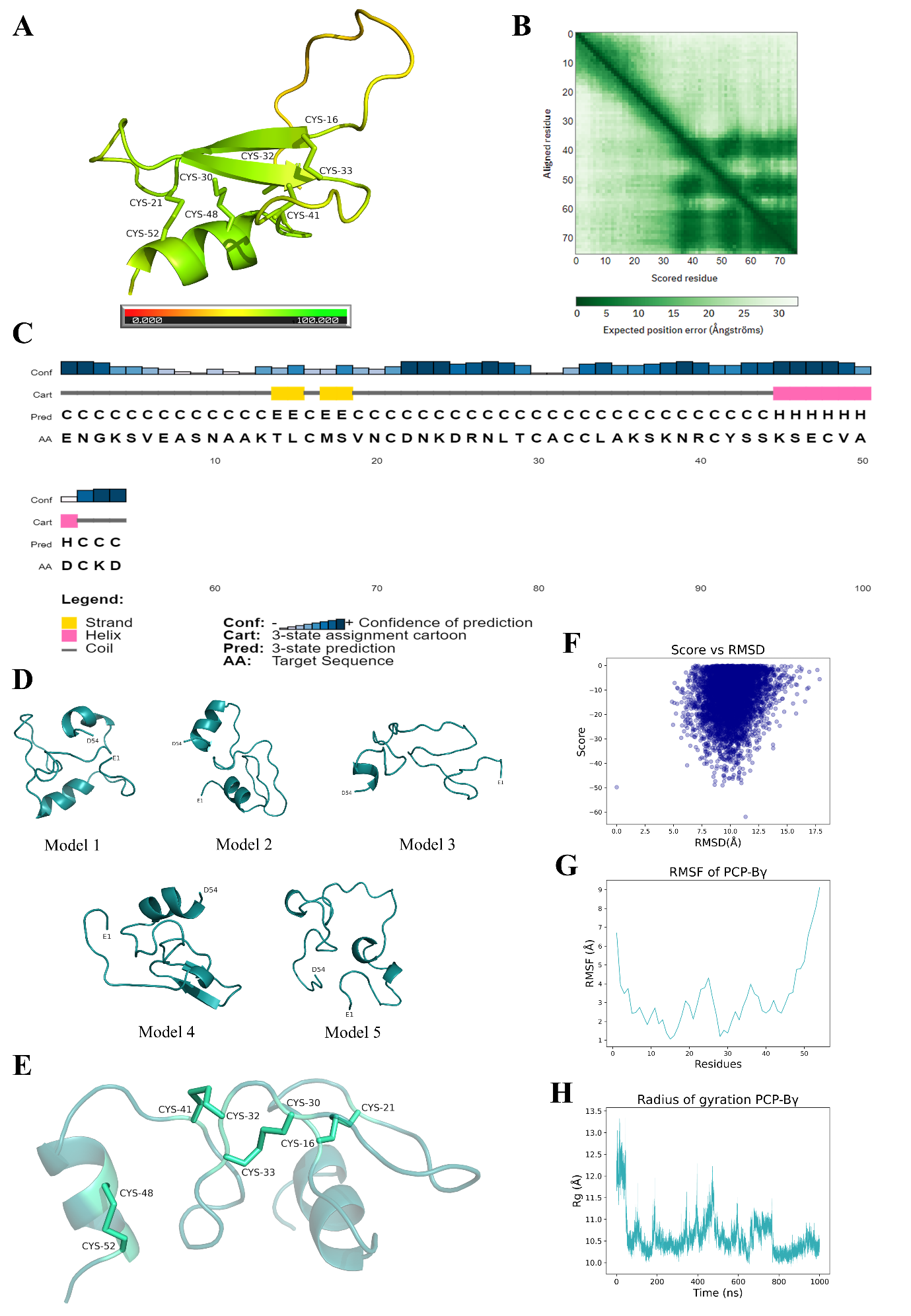

**Supplementary Figure S2.**

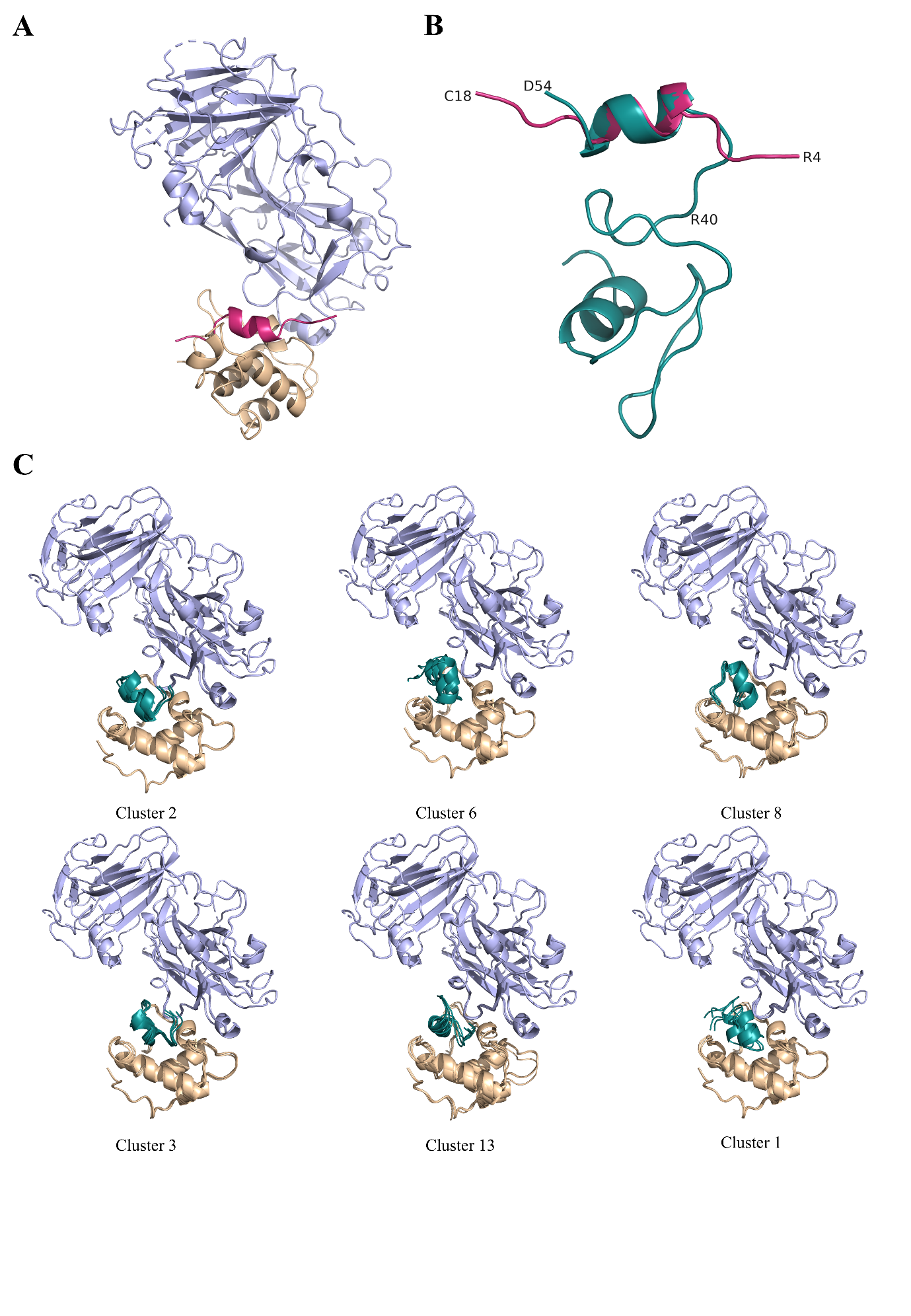

**Supplementary Figure S3.**

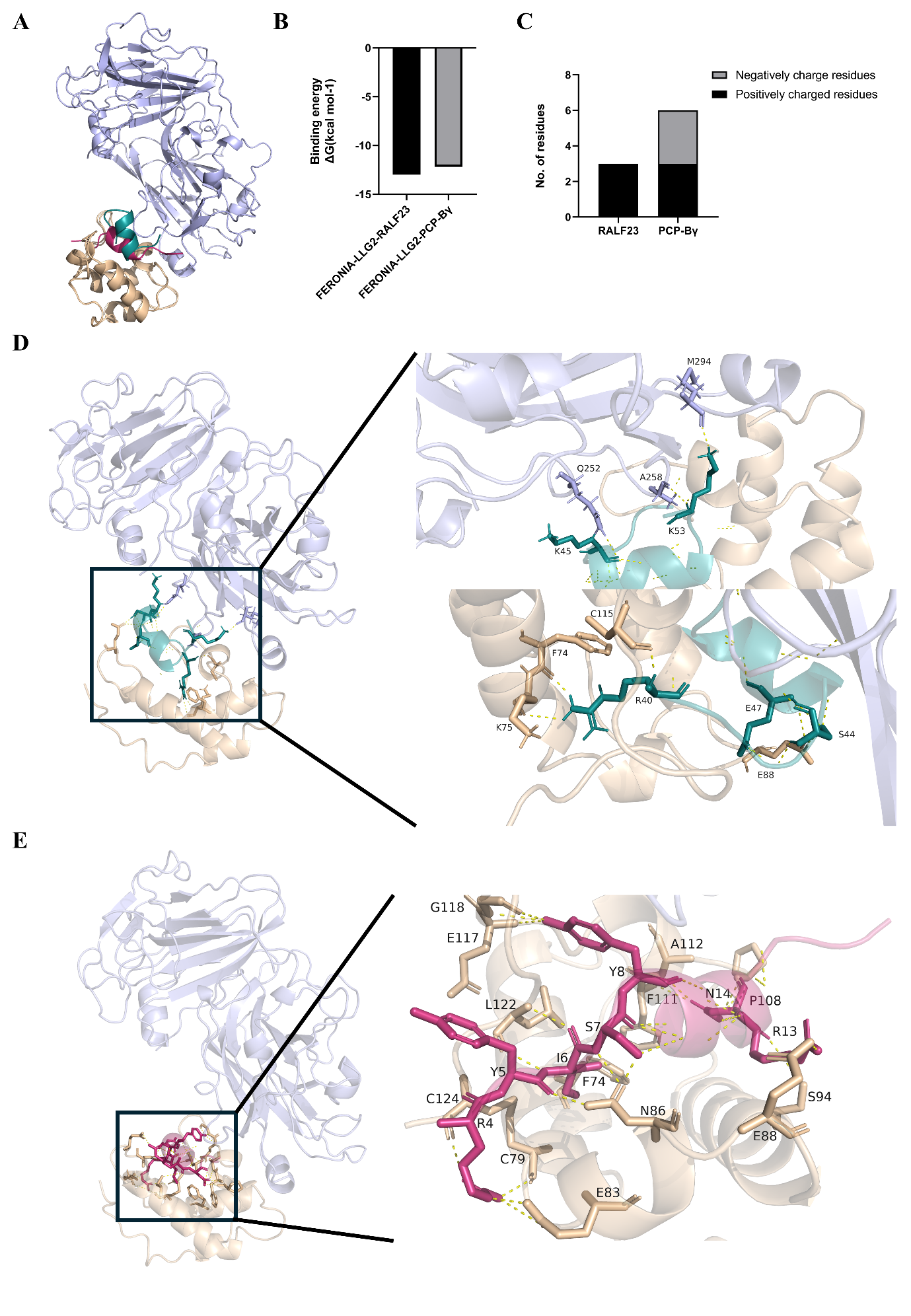

**Supplementary Figure S4.**

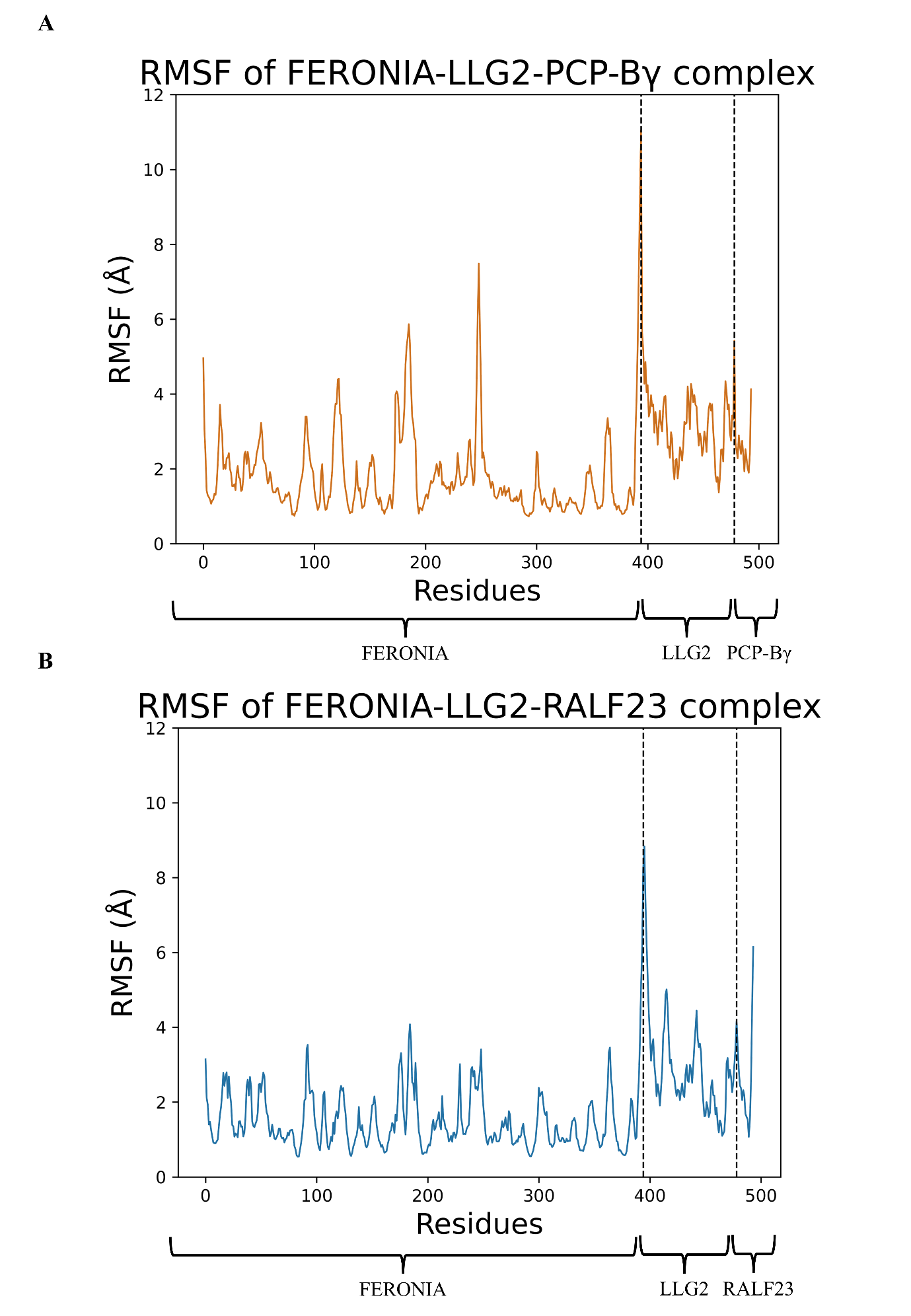

**Supplementary Figure S5.**

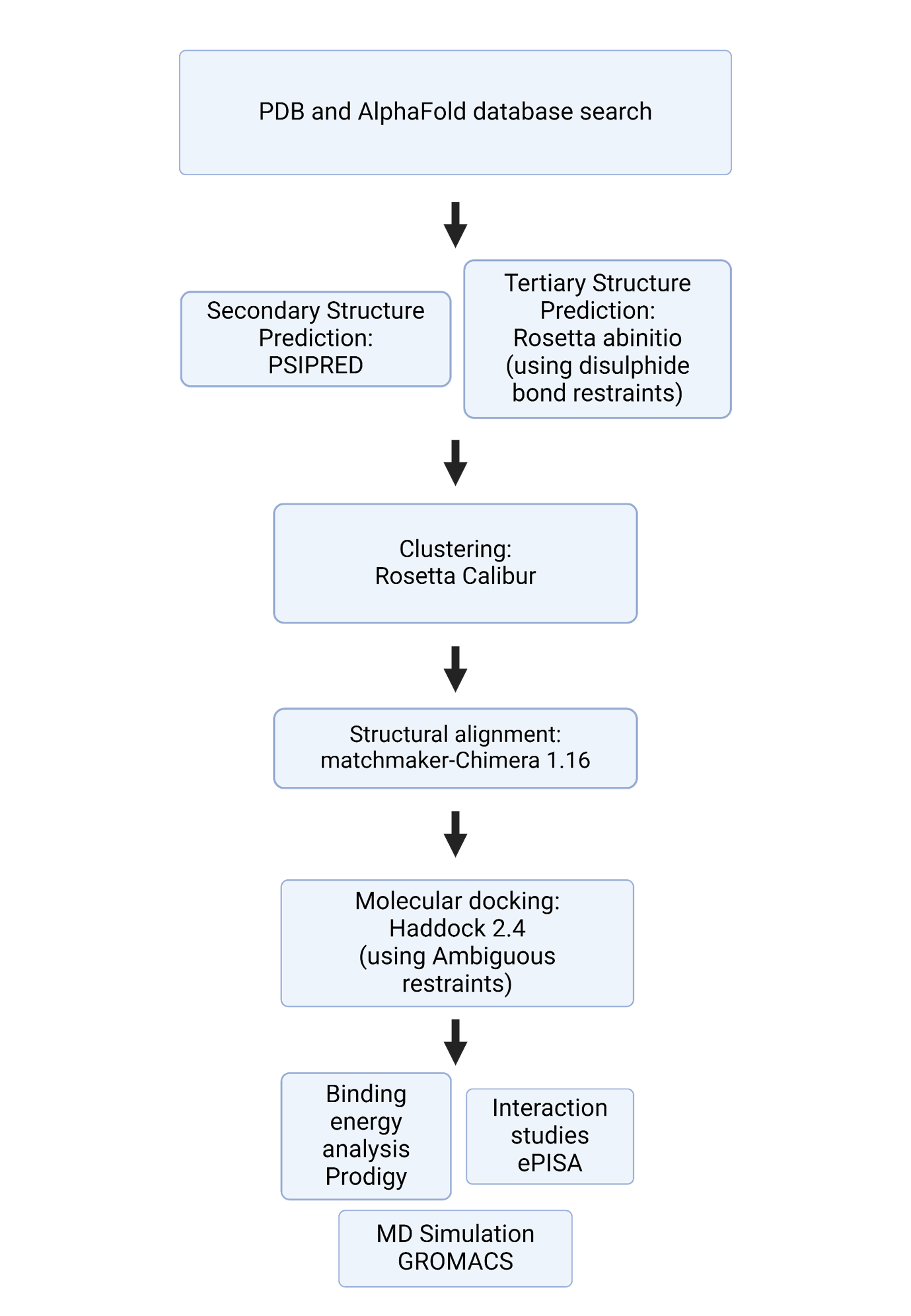

**Supplementary Tables:**

**Supplementary Table S1. Disulfide bond structural restraints used during PCP-Bγ modeling using Rosetta abinitio protocol.**

| Cysteine position | Residue number | Cysteine position | Residue number |
| --- | --- | --- | --- |
| Cys1 | 16 | Cys2 | 21 |
| Cys3 | 30 | Cys5 | 33 |
| Cys4 | 32 | Cys6 | 41 |
| Cys7 | 48 | Cys8 | 52 |

**Supplementary Table S2. Cluster size of predicted PCP-Bγ determined using Calibur.**

| Cluster | Size |
| --- | --- |
| Cluster 1 | 1802 |
| Cluster 2 | 1152 |

**Supplementary Table S3. Top 5 scoring models of** **PCP-Bγ predicted using ROSETTA abinitio protocol.**

***Model used for further analysis**

| Model No. | ROSETTA Score | RMSD |
| --- | --- | --- |
| 1 | -61.953 | 11.33 |
| 2* | -49.834 | 0 |
| 3 | -49.062 | 9.305 |
| 4 | -48.840 | 8.613 |
| 5 | -48.123 | 10.970 |

**Supplementary Table S4. Ambiguous restraints provided during molecular docking.**

| Molecule | Residue Number | Molecule | Residue Number |
| --- | --- | --- | --- |
| FERONIA | 256  257  258  260  261  262  295  296 | PCP-Bγ | 40  41  42  43  44  45  46  47  48  49  50  51  52  53  54 |
| LLG2 | 75  79  82  83  85  86  88  93  94  97  101  108  109  112  115  116  117  118  119  120  121  122  123  124  125 | PCP-Bγ | 40  41  42  43  44  45  46  47  48  49  50  51  52  53  54 |

**Supplementary Table S5. Predicted FERONIA-PCP-Bγ-LLG2 clusters by Haddock 2.4, their respective haddock score, cluster size, and root mean square deviation (RMSD). The top 6 clusters are highlighted in bold.**

***Cluster used for further analysis**

| Cluster | Cluster Size | Haddock score | RMSD(Å) |
| --- | --- | --- | --- |
| **1** | **30** | **-58.282** | **0.409** |
| **2*** | **28** | **-82.303** | **0.226** |
| **3** | **26** | **-65.917** | **0.251** |
| 4 | 20 | -39.972 | 0.552 |
| 5 | 14 | -46.467 | 0.398 |
| **6** | **11** | **-76.040** | **0.571** |
| 7 | 7 | -37.974 | 0.324 |
| **8** | **7** | **-71.619** | **0.296** |
| 9 | 6 | -42.154 | 0.756 |
| 10 | 5 | -45.915 | 0.558 |
| 11 | 4 | -31.034 | 0.608 |
| 12 | 4 | -43.937 | 0.215 |
| **13** | **4** | **-58.700** | **0.390** |

**Supplementary Table S6. Binding energies predicted using Prodigy for FERONIA-LLG2-RALF23 crystal structure, and FERONIA-LLG2-PCP-Bγ predicted model.**

| Model | Binding energy  ΔG (kcal mol^-1^) | K_d_  (M) at 25˚C |
| --- | --- | --- |
| FERONIA-LLG2-RALF23 | -13.0 | 3.1 x 10^-10^ |
| FERONIA-LLG2-PCP-Bγ | -12.2 | 1.1 x 10^-09^ |

**Supplementary Table S7. Polar contacts between various residues of FERONIA-LLG2-PCP-Bγ predicted model.**

| Molecule | Residue | Molecule | Residue |
| --- | --- | --- | --- |
| PCP-Bγ | K53 | FER | A258  M294 |
|  | K45 |  | Q252 |
|  | E47 | LLG2 | E88 |
|  | R40 |  | C115  F74  K75 |

**Supplementary Table S8. Polar contacts between various residues of FERONIA-LLG2-RALF23 crystal structure.**

| Molecule | Residue | Molecule | Residue |
| --- | --- | --- | --- |
| RALF23 | R4 | LLG2 | C124  E83  C79 |
|  | Y5 |  | N86 |
|  | I6 |  | L122 |
|  | S7 |  | N86 |
|  | Y9 |  | E117  G118 |
|  | R13 |  | S94  E88 |
|  | N14 |  | P108  F111  A112 |
